# Domain shifts in infant brain diffusion MRI: Quantification and mitigation for fODF estimation

**DOI:** 10.1101/2025.04.04.647303

**Authors:** Rizhong Lin, Hamza Kebiri, Ali Gholipour, Yufei Chen, Jean-Philippe Thiran, Davood Karimi, Meritxell Bach Cuadra

## Abstract

The accurate estimation of fiber orientation distributions (fODFs) in diffusion magnetic resonance imaging (MRI) is crucial for understanding early brain development and its potential disruptions. While deep learning (DL) models have shown promise in fODF estimation from neonatal diffusion MRI (dMRI) data, the out-of-domain (OOD) performance of these models remains largely unexplored, especially under diverse domain shift scenarios. This study evaluated the robustness of three state-of-the-art DL architectures—MLP-based, Transformer-based, and U-Net/CNN-based—on fODF predictions derived from diffusion MRI data. Using 488 subjects from the developing Human Connectome Project (dHCP) and the Baby Connectome Project (BCP) datasets, we reconstructed reference fODFs from the full dMRI series using single-shell three-tissue constrained spherical deconvolution (SS3T-CSD) and multi-shell multi-tissue CSD (MSMT-CSD) to generate reference fODF reconstructions for model training, and systematically assessed the impact of age, scanner/protocol differences, and input dimensionality on model performance. Our findings reveal that U-Net consistently outperformed other models when fewer diffusion gradient directions were used, particularly with the SS3T-CSD-derived ground truth, which showed superior performance in capturing crossing fibers. However, as the number of input diffusion gradient directions increased, MLP and the transformer-based model exhibited steady gains in accuracy. Nevertheless, performance nearly plateaued from 28 to 45 input directions in all models. Age-related domain shifts were found to be less pronounced in late developmental stages (late neonates, and babies), with SS3T-CSD demonstrating greater robustness to variability compared to MSMT-CSD. To address inter-site domain shifts, we implemented two adaptation strategies: the Method of Moments (MoM) and fine-tuning. Fine-tuning consistently yielded superior results, with U-Net benefiting the most from increased target subjects. This study represents the first systematic evaluation of OOD settings in deep learning applications to fODF estimation, providing critical insights into model robustness and adaptation strategies for diverse clinical and research applications.

## 1 Introduction

Early brain development is a crucial period that sets the stage for lifelong health (Bhat et al., 2014; Godfrey and Barker, 2001; O’Donnell and Meaney, 2017; Volpe, 2009). The depiction of white matter fiber bundles, which are responsible for relaying action potential signals between different brain areas, is of particular interest for fetal, newborn, and baby brains. Those long and myelinated axons have been shown to play a significant role in cognitive and motor functions from infancy (Dubois et al., 2014) to adulthood (Brun and Englund, 1986; Davis et al., 2003; Ruiz-Rizzo et al., 2024). Precise estimation of these bundles of fibers is essential to comprehend *in-vivo* developmental trends and identify irregularities that might indicate potential diseases.

Progress in diffusion magnetic resonance imaging (dMRI), a non-invasive technique that relies on water molecule displacement as a proxy to microstructure, has yielded unparalleled insights into the mapping of the human brain (Descoteaux et al., 2011; Özarslan et al., 2013). The predominant method for extracting diffusion properties from the diffusion signal typically involves a prior model, commonly the diffusion tensor imaging (DTI) model (Basser et al., 1994). However, more intricate models such as the multi-shell multi-tissue constrained spherical deconvolution (MSMT-CSD) aim to reconstruct Fiber Orientation Distribution Functions (fODFs), enabling the representation of complex white matter configurations like fiber crossings (Jeurissen et al., 2014a; Tournier et al., 2004) with sufficiently large crossing angles (Schilling et al., 2018). These models, however, necessitate densely sampled multi-shell dMRI data that require high acquisition times. A less-data demanding method has been recently proposed: single-shell three-tissue constrained spherical deconvolution (SS3T-CSD) (Dhollander and Connelly, 2016; Dhollander et al., 2016) reconstructs multi-tissue data with a single non-zero b-value and was demonstrated to be a good fit for developing brains (Dhollander et al., 2019) in which white matter voxels might suffer more from partial volume effect.

### 1.1 Estimating fODFs with machine learning

With machine and more prominently deep learning, it has become possible to efficiently estimate the mapping between different related diffusion quantities, for instance from dMRI signals to scalars such as fractional anisotropy (Alexander et al., 2014; Golkov et al., 2016; Karimi and Gholipour, 2022; Tian et al., 2020). Also, fODF prediction from the original raw signal or its spherical harmonic representation has raised a growing interest from the community in recent years (Bartlett et al., 2023; Hosseini et al., 2022; Jha et al., 2023; Karimi et al., 2021b; Kebiri et al., 2023b; Lin et al., 2019; Nath et al., 2019c). Specifically, Nath et al. (2019c) used a dense residual network to predict fODFs derived from *ex-vivo* confocal microscopy images of monkey histology sections. However, this method is constrained by the unavailability of *ex-vivo* histological training data. While Karimi et al. (2021a) used a voxel-wise multilayer perceptron (MLP) to predict fODFs, Bartlett et al. (2023); Lin et al. (2019) employed a 3D convolutional neural network (CNN) to predict fODFs of the central voxel based on a small neighborhood of the diffusion signal. To further exploit the correlations among neighboring voxels, a two-stage Transformer-CNN was employed by Hosseini et al. (2022) to convert 200 measurements into 60 measurements before proceeding to predict fODFs. Another work Jha et al. (2023) has shown, using a differential equation approach, the feasibility of predicting accurate fODFs with a limited number of diffusion gradient directions using a 2D neighborhood. On the other hand, Kebiri et al. (2023b) used a 3D U-Net-like network with extensive residual connections to predict big patches, leveraging spatial correlations to estimate fODFs with a small number of input measurements and hence a substantial reduction in scanning times. This approach has yielded promising results on newborns and fetuses (Kebiri et al., 2024).

Differently, Koppers and Merhof (2016) have tried to estimate the orientation of the fibers using 2D CNN in a classification paradigm. Other studies have aimed at segmenting fiber tracts either through a prior model applied on the input (Dong et al., 2019; Wasserthal et al., 2018) or directly from a spherical representation of the acquired signal (Kebiri et al., 2023a). In a different paradigm, da Silva et al. (2024) have successfully used deep learning to map an SS3T-CSD fODF constructed with a single b-value acquisition with a limited number of directions to an MSMT-CSD fODF reconstructed with a multi-shell high angular acquisition. An extensive review of machine learning applications in diffusion MRI can be found in Karimi and Warfield (2024); Kebiri (2023).

### 1.2 Domain shifts in dMRI and mitigation strategies

While DL applied in medical imaging offers strong advantages as detailed above, it suffers substantially from *domain shift* issues in which source and target data distributions vary considerably. In fact, small datasets that are limited in age span and the privacy constraints of sharing them at scale hamper cross-dataset studies. In MRI (Richiardi et al., 2025), domain shift is even more amplified as scanners from different sites vary in brands and field strengths and sequence acquisition parameters that differ significantly, even within one modality such as dMRI (b-values and the gradient directions as an example). Both biological shifts (Bento et al., 2022; Dubois et al., 2014), such as age or pathology, and technological shifts (Tax et al., 2019) contribute to the final distribution shift between the source sets and the target sets. Age is a particular shift in developing brains because of the rapid change in structure and function (Konkel, 2018; Schilling et al., 2023).

To mitigate distribution shifts in MRI, solutions may operate at the data level (e.g., data harmonization or augmentation) or at the model level (e.g., transfer learning or domain adaptation).

Data harmonization, which aims to minimize differences due to the unwanted shift between the source domain and the target domain, is dominated by statistical methods (Huynh et al., 2019; Johnson et al., 2007; Karayumak et al., 2019; Karimi and Warfield, 2024; Mirzaalian et al., 2018). The most dominant ones are the Rotation Invariant Spherical Harmonics (RISH) (Karayumak et al., 2019; Mirzaalian et al., 2018), specifically designed for the original dMRI signal, and COMBAT (Johnson et al., 2007), a more general harmonization method that is applied to the target diffusion map (Pinto et al., 2020), i.e. after model fitting. The Method of Moments (MoM) (Huynh et al., 2019), which aligns diffusion-weighted imaging (DWI) features via spherical moments, was also proposed recently and achieved promising results in developing brains (Lin et al., 2024a). While most harmonization methods, such as RISH, require similar acquisition protocols and site-matched healthy controls, MoM and Combat are not subject to these restrictions, making them appropriate to a broader range of conditions.

Deep learning has been also used for harmonizing dMRI metrics (Hansen et al., 2022; Koppers et al., 2019; Moyer et al., 2020; Nath et al., 2019b), including those for fODF estimation. However, they either need paired acquisitions with histology (Nath et al., 2019a,b) or scan-rescan (Yao et al., 2023b,c) acquisitions. Although deep learning techniques offer solutions to nonlinear harmonization, they are prone to overfitting and require extensive training data, often from matched acquisitions, that are not easy to get (Bashyam et al., 2022; Pinto et al., 2020).

Domain adaptation methods have been employed to tackle domain shifts in medical imaging (Guan and Liu, 2022). However, only two methods have been proposed in the context of dMRI, and they aim to tackle the diversity of dMRI acquisitions and, in particular, the b-value. Kamphenkel et al. (2018) circumvented the domain shift by using a diffusion kurtosis model to estimate missing input values in a breast cancer classification task, while Yao et al. (2023a) used a dynamic head to learn the different shell configurations using spherical convolutions to predict fODFs. However, these methods do not offer robustness to other shifts such as scanner, age, or different protocols (except the b-value). Using deep learning, two orthogonal approaches are particularly interesting: adversarial training and transfer learning (fine-tuning). The former relies on learning invariant features through a domain-agnostic loss function (Ganin et al., 2016; Kamnitsas et al., 2017) while the latter relies on pre-trained weights from a target-related dataset (Ghafoorian et al., 2017; Samala et al., 2018), that can range from a source dataset to a public dataset of natural images (Krizhevsky et al., 2012).

### 1.3 Contributions of this work

The exploration of fODF estimation under domain shifts remains limited. While Karimi et al. (2021b) and Kebiri et al. (2023b) have extensively tested their fODF prediction models on developing neonatal brains, only Kebiri et al. (2023b) has investigated out-of-domain (OOD) performance. However, this OOD evaluation was conducted qualitatively due to the lack of fetal fODF ground truth. Our preliminary work in Lin et al. (2024a) explored the age- and age/scanner/protocol-related domain shifts and proposed potential solutions, including the Method of Moments (data harmonization) and fine-tuning (domain adaptation).

In this study, we significantly extend (Lin et al., 2024a) by including three state-of-the-art deep learning models, MLP-based, Transformers-based, and U-Net/CNN-based. We use also two cohorts: the newborns of the developing Human Connectome Project (dHCP) and the babies of the Baby Connectome Project (BCP). We extend our analysis to different fODF *ground-truth* models: SS3T-CSD (Dhollander and Connelly, 2016; Dhollander et al., 2016) and MSMT-CSD (Jeurissen et al., 2014a). Furthermore, different methodological configurations are evaluated: the number of input diffusion gradient directions to the model and the number of subjects from the target set used for domain adaptation/harmonization. To the best of our knowledge, this is the first study to assess OOD settings of DL applications to fODFs estimation thoroughly.

## 2 Methods

Our framework for fODF prediction and OOD evaluation is illustrated in Figure 1 in multiple stages. First, reference Constrained Spherical Deconvolution (CSD) algorithms 𝒩 ∈{(MSMT-CSD, SS3T-CSD)} generate ground truth fODFs from full diffusion MRI series for training. Next, deep learning models 𝒩∈ MLP, CTtrack, U-Net undergo supervised training using source domain data (*S*_*src*_) to predict these reference fODFs. The framework is evaluated in both intra-site and inter-site scenarios (detailed in Section 3), with domain shifts addressed through either data harmonization using Method of Moments to transform target domain data (*S*_*tar*_) before inference, or model adaptation via fine-tuning on target domain data to create adapted networks (𝒩 ^′^).

**Figure 1.**
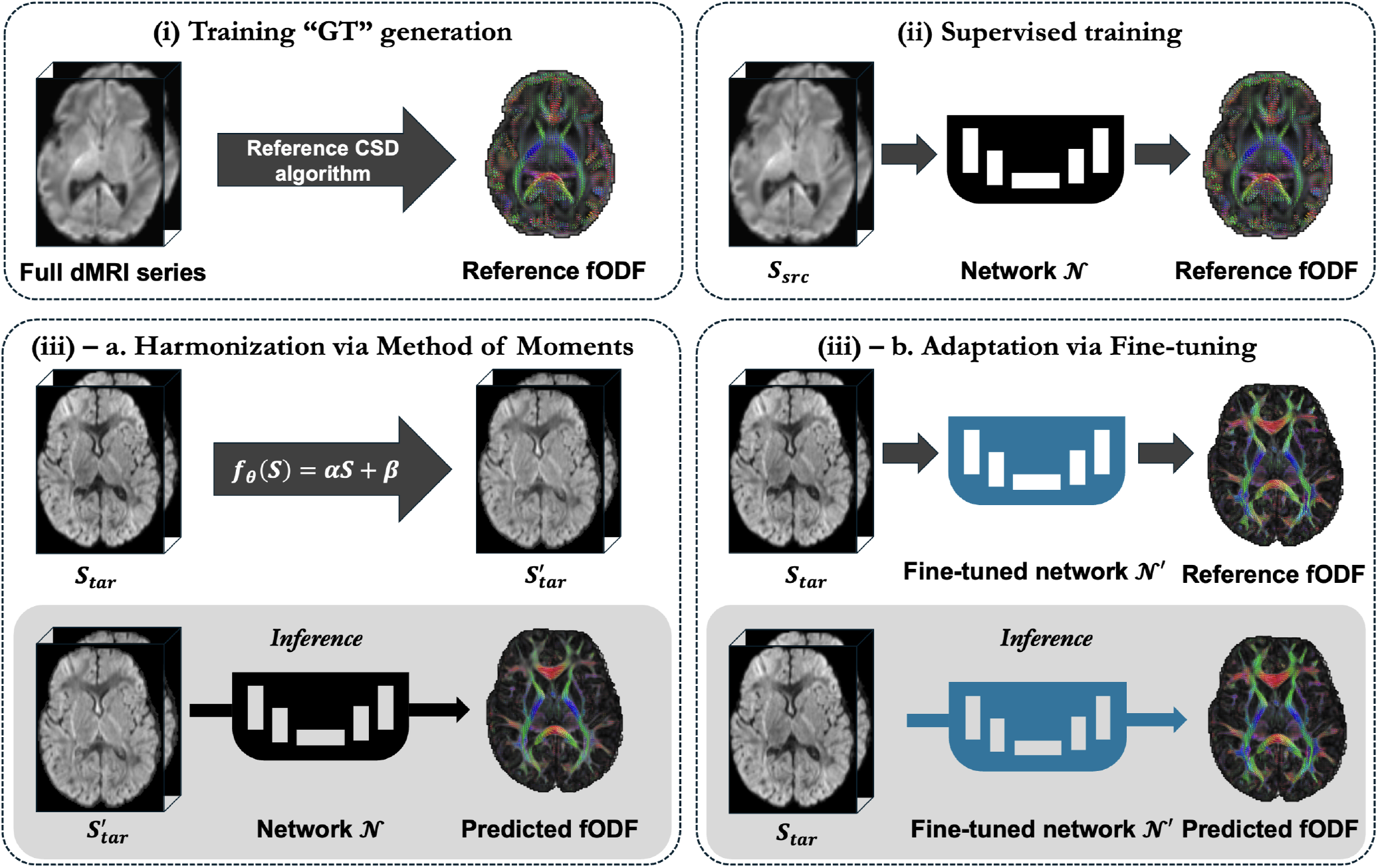
Overview of the fODF prediction framework using multiple deep learning architectures 𝒩 ∈ MLP, CTtrack, U-Net. (i) Reference fODFs are generated from full dMRI series using CSD algorithms 𝒩 ∈ {(MSMT-CSD or SS3T-CSD)} as ground truth for training. (ii) Models are trained on source domain data to predict these reference fODFs. (iii) Domain shifts (both intra-site and inter-site) are addressed through either: a) Method of Moments harmonization to transform target data before inference with the original model, or b) Fine-tuning the model on target domain data to create an adapted network. The framework enables comprehensive evaluation across different acquisition protocols and age ranges, as detailed in Section 3.

### 2.1 Backbone model architectures

Three distinct deep learning architectures were implemented for fODF prediction, each representing different modeling paradigms: a U-Net-based architecture for spatial context learning (Kebiri et al., 2023b), a hybrid CNN-Transformer for local-global feature integration (Hosseini et al., 2022), and an MLP for direct signal-to-fODF mapping (Karimi et al., 2021b). These architectures share a common input-output framework: input diffusion measurements are normalized and projected onto a spherical harmonics (SH) basis to enhance acquisition independence, with SH order selection determined by the number of input directions. The output consistently represents fODF in SH basis (SH-*L*_max_ order 8) with 45 coefficients, denoted as *n*_sh_ = 45.

#### 2.1.1 U-Net based method

The network architecture extends U-Net (Ronneberger et al., 2015) with extensive short and long-range residual connections and stride-2 convolutions replacing conventional max-pooling operations. The network processes patches of 16×16×16× *n*_sig_ voxels and outputs corresponding fODF 16×16×16×45. The first block contains 36 feature maps, doubled after each contracting block. Convolutions are followed by ReLU activation (Agarap, 2018) and dropout (Srivastava et al., 2014) layers, except in the output layer.

#### 2.1.2 CNN-Transformer (CTtrack)

This hybrid architecture uses 3D CNNs for initial feature extraction, followed by a Transformer network (Hosseini et al., 2022). A×3×3×3×*n*_sig_ patch is processed by CNN layers, producing feature embeddings for each voxel. These embeddings, combined with positional encodings, are fed into four self-attention layers. An MLP head processes the aggregated features to estimate 45 SH coefficients for the central voxel.

#### 2.1.3 Multilayer perceptron (MLP)

The MLP architecture (Karimi et al., 2021b) comprises an input layer, six hidden layers, and an output layer with neuron configuration [*n*_sig_, 300, 300, 300, 400, 500, 600, *n*_sh_]. Each hidden layer can include nonlinear activations and dropouts. The output layer predicts *n*_sh_ = 45 SH coefficients for the fODF.

### 2.2 CSD-based fODF reconstruction algorithms for training

To generate *training ground truths*, we used two classical fODF estimation models, Multi-Shell Multi-Tissue CSD (MSMT-CSD) (Jeurissen et al., 2014b) and Single-Shell 3-Tissue CSD (SS3T-CSD) (Dhollander and Connelly, 2016) under various dMRI acquisition settings. Following established terminology (Lin et al., 2024b), we refer to these fODF reconstructions as “ground truth” or “GT” when used as training targets for the deep learning models, while acknowledging these are reference reconstructions rather than true anatomical measurements. MSMT-CSD leverages multi-shell data to compute tissue-specific response functions and yields multi-tissue fODFs. SS3T-CSD uses single-shell (+b0) data and a three-tissue approach, enabling approximate multi-tissue decomposition with reduced acquisition demands. These reconstructions from classical models serve as training targets for the deep learning models.

### 2.3 Training details

The U-Net training employed Adam optimizer (Kingma and Ba, 2014) with *ℓ*_2_ norm loss, initial learning rate 2×10^−4^, 0.1 dropout rate, and batch size 1, processing 128 patches per epoch for 500-1000 epochs with early stopping. CTtrack training used AdamW optimizer (Loshchilov and Hutter, 2019) with initial learning rate 1×10^−3^, weight decay 5×10^−3^, batch size 4000, and *ℓ*_1_ loss function for 100 epochs. The MLP employed Adam optimizer with initial learning rate 1×10^−3^, batch size 2000, and *ℓ*_2_ norm loss for 100 epochs. Convergence times approximated 12, 12, and 4 hours for U-Net, CTtrack, and MLP, respectively, with inference requiring less than 10 seconds per subject.

### 2.4 Domain shift mitigation methods

#### 2.4.1 Data harmonization based on Method of Moments

We used the Method of Moments (MoM) approach (Huynh et al., 2019) to harmonize diffusion MRI data across different sites or acquisition protocols, as it is not subject to protocol or subject matched constraints. A linear transformation *f*_*θ*_ (*S*) = *αS*+ *β* matches the statistical moments of source and target datasets. For each gradient direction, we compute the mean (*m*_1_) and variance (*m*_2_) of the target and reference data within brain masks. Voxel-wise estimates of *α* and *β* are obtained by minimizing

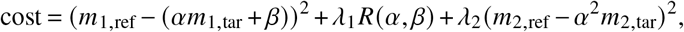

where *R*(*α, β*) is a regularization term penalizing large deviations from identity:

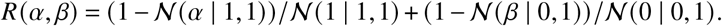

#### 2.4.2 Domain adaptation by fine-tuning

The domain adaptation framework employs transfer learning through model fine-tuning, maintaining the same architecture while adjusting network parameters to target domain distribution.

Each model maintained its original optimizer while adapting to target domain data. The U-Net processed target data in 16^3^ patches with stride 8 voxels (50% overlap in each dimension), using only patches within the brain mask. These patches were split into training and validation sets (4:1 ratio). Fine-tuning ran for 20 epochs with learning rate 1× 10^−5^ and batch size 64 for U-Net, while CTtrack and MLP used 10 epochs with learning rates 1× 10^−4^ and 2× 10^−4^ respectively, and batch size 2000.

### 2.5 Implementation details

All models were implemented using PyTorch (Paszke et al., 2019), PyTorch Lightning (Falcon and The PyTorch Lightning team, 2024), MONAI (Cardoso et al., 2022), and TensorFlow 2 (Abadi et al., 2015). The U-Net architecture was reimplemented from its original Tensorflow version using PyTorch. CTtrack maintained its original TensorFlow 2 implementation with modified data loading pipelines. The MLP architecture was reconstructed in TensorFlow 2 following published specifications. The method of moments harmonization was implemented in MATLAB R2022b. Training used an NVIDIA GeForce RTX 2080 Ti with 11GB RAM and 24 CPU cores.

### 2.6 Evaluation

A quantitative assessment was carried out to evaluate the performance of the fODF estimation methods. The evaluation process relied on three metrics: (1) the agreement in terms of the number of peaks quantifying the rate of concordance between two methods in the number of fibers predicted, (2) mean angular error between those peaks, and (3) the apparent fiber density (AFD) (Raffelt et al., 2012), defined as the amplitude of the fODF. The peaks were computed using Dipy (Garyfallidis et al., 2014) with a mean separation angle of 45°, a maximum of 3 peaks, and a relative peak threshold of 0.5. Details about these metrics can be found in Kebiri et al. (2024). The AFD error/difference is computed as a masked mean absolute percentage error (MAPE), where the mask is confined to white matter voxels, and is given by: 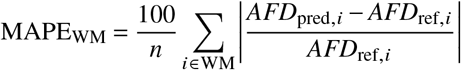, where *n* is the number of white matter voxels.

## 3 Experiments

The proposed framework was extensively evaluated through a series of experiments designed to assess both the performance of different network architectures 𝒩 ∈ {( MLP, CTtrack, U-Net)} and their robustness to various domain shifts. We first validate the CSD-based reconstruction methods used for ground truth generation, then investigate intra-site domain shifts (particularly age-related variations), and finally examine inter-site domain shifts between two major developing brain imaging consortia. The effectiveness of our domain shift mitigation strategies is evaluated across these scenarios.

### 3.1 Datasets and preprocessing

#### 3.1.1 Neonates of the developing Human Connectome Project (dHCP)

We use the third data release of the publicly available dHCP dataset^1^ (Edwards et al., 2022) which was acquired on a 3T Philips Achieva system with a 32-channel neonatal head coil. The protocol employed TE = 90 ms, TR = 3800 ms, a multiband factor of 4, a SENSE factor of 1.2, a Partial Fourier factor of 0.855, in-plane resolution 1.5 mm^2^, slice thickness 3 mm with 1.5 mm overlap (Hutter et al., 2018), and four shells {0, 400, 1000, 2600} s/mm^2^ with 20, 64, 88, and 128 volumes. Data were preprocessed with SHARD (Christiaens et al., 2021; Pietsch et al., 2021), including MP-PCA denoising (Veraart et al., 2016), motion and distortion correction, Gibbs suppression, and resampling to 1.5 mm^3^ isotropic resolution of size 100× 100× 64 voxels. White matter masks were obtained by combining T2-weighted segmentation labels *White Matter* and *Brainstem* (registered to diffusion space (Yushkevich et al., 2016)) with FA > 0.25 computed in MRtrix (Tournier et al., 2012), following Kebiri et al. (2024). We used a total of 323 unique subjects from dHCP. Of these, 165 subjects (PMA [postmenstrual age] at scan: [35.57, 45.29] weeks, median: 40.29, mean: 40.01, SD: 2.29) were denoted as dataset *S*_dHCP_. Additionally, we selected two age-distinct sets from dHCP, denoted as *S*_dHCP,early_ and *S*_dHCP,late_ respectively, each consisting of 105 subjects: early-stage (PMA at scan: [33.14, 37.86] weeks, median: 35.57, mean: 35.65, SD: 1.39) and late-stage (PMA at scan: [41.0, 45.14] weeks, median: 42.43, mean: 42.48, SD: 1.07).

#### 3.1.2 Babies of the Human Connectome Project (BCP)

We used the data from the publicly available Baby Connectome Project (BCP) dataset^2^ (Howell et al., 2019). Images were acquired on a 3T Siemens Magnetom Prisma with a 32-channel head coil. The dMRI protocol used six shells *b* ∈{500, 1000, 1500, 2000, 2500, 3000} s/mm^2^ having 9, 12, 17, 24, 34, 48 diffusion gradient directions respectively, and 6 *b0* images. Other parameters included TE = 88.6 ms, TR = 2640 ms, multiband factor 5, resolution 1.5 mm^3^, with a field of view of 140× 140× 96 voxels. Preprocessing included denoising, bias correction, motion compensation, and distortion correction (Andersson and Sotiropoulos, 2016), followed by FSL BET brain extraction (Jenkinson et al., 2012). White matter masks were derived by combining a SynthSeg^+^ (Billot et al., 2023) WM mask, voxels with FA > 0.4, and voxels with (FA > 0.15) ∧ (MD > 0.0011). We used 165 subjects from BCP (age at scan: [2.0, 24.0] months, median: 15.00, mean: 15.91, SD: 5.87) and denote this dataset as *S*_BCP_.

### 3.2 Validation of CSD-based reconstruction methods

To assess the quality and consistency of the reconstruction methods used to generate training targets, we split each subject’s diffusion MRI series into two equal, non-overlapping subsets (GS1 and GS2), following Kebiri et al. (2023b). For the dHCP dataset, each subset contained 150 measurements distributed across b-values {0, 400, 1000, 2600} s/mm^2^ with 10, 32, 44, and 64 directions, respectively. Similarly, for the BCP dataset, each subset contained 75 measurements. We reconstructed fODFs using MSMT-CSD on both datasets using all available b-values. For the dHCP dataset, we additionally performed SS3T-CSD reconstruction using only the b = 0 and b = 1000 s/mm^2^ shells.

Since GS1 and GS2 represent equivalent samplings of the same underlying diffusion signal, the difference between their reconstructed fODFs indicates the inherent variability in each reconstruction method. This comparison serves as an upper bound for the expected error between the learning-based fODF estimation methods and their training targets, which are generated by applying these reconstruction methods on the complete diffusion MRI series. For the dHCP dataset, we also compared the fODFs reconstructed from the complete series (300 measurements for MSMT-CSD, and b0/b1000 measurements for SS3T-CSD) to assess and understand the differences between these two reconstruction approaches.

### 3.3 Intra-site domain shift quantification

We evaluated within-domain performance using both *S*_dHCP_ and *S*_BCP_ datasets as defined in Section 3.1, each containing 165 subjects. The subjects were split into 85/80 for training-validation/testing. Age-related variations were investigated exclusively in the dHCP dataset, as BCP data showed minimal age-related variation (Lin et al., 2024a). For age-specific analysis, we used *S*_dHCP,early_ and *S*_dHCP,late_, containing 105 subjects each, split 50/55 for training-validation/testing.

For input signals, we used varying numbers of directions (*n*_sig_): {6, 15, 28, 45} for dHCP and {6, 12} for BCP, with normalization by b0. Ground truth fODFs were generated using MSMT-CSD for both datasets. For dHCP, we conducted additional experiments using SS3T-CSD-generated fODFs from all 88 b1000 and 20 b0 measurements as an alternative training ground truth. For baseline comparisons, we performed in-domain testing (training and testing on the same age group).

### 3.4 Inter-site domain shift quantification

This analysis quantifies performance degradation when training on *S*_dHCP_ but testing on *S*_BCP_ and vice versa. The mismatch includes different MRI scanners (Philips 3T for dHCP, Siemens 3T for BCP), acquisition protocols, and subject age ranges. We used 165 subjects from each dataset (*S*_dHCP_ and *S*_BCP_), split into 85/80 for training-validation/testing. Both datasets were normalized by *b0*, and we investigated *n*_sig_ ∈ {6, 12} for each network, comparing how domain shift impacts the fODF estimation results.

### 3.5 Domain shift attenuation experiments

We evaluated two domain shift mitigation strategies when transferring models between the two sites (dHCP and BCP), as described in Section 2.4. Using {1, 2, 5, 10} target subjects, we tested both Method of Moments (MoM) harmonization, which transformed the input dMRI signals to match target domain statistics, and model fine-tuning, which utilized both dMRI signals and ground truth fODFs to adapt the pre-trained source model. As shown in Figure 1, we compared two inference scenarios: (1) applying the original model to harmonized target data and (2) using the fine-tuned model on the original target data. These experiments quantified the effectiveness of each approach in improving fODF reconstruction quality across domains.

## 4 Results

### 4.1 Ground-truth reconstruction consistency

Table 1 shows the consistency between fODFs reconstructed from GS1 and GS2 of each subject using classical reconstruction methods. Single-fiber estimations show a good agreement rate (92%) for dHCP and 86% for BCP) for MSMT-CSD, but a low agreement is reported for both datasets of around 45% and 16° in angular error. SS3T-CSD exhibits higher agreement for multiple fibers (56%) and lower for single fibers (63%). As reported previously Lin et al. (2024b), the proportion of multiple fibers predicted by SS3T-CSD (61%) is closer to literature values (66%) compared to MSMT-CSD (23%). Moreover, the number of b1000 in the dHCP dataset is around half of the minimum required to estimate the 45 coefficients of the SS3T fODFs. Hence, more data will likely result in higher agreement for single and multiple fibers.

**Table 1.** Consistency analysis between split-dataset fODF reconstructions (ΔGS). Agreement rates and angular differences are shown for single and two-fiber configurations, along with AFD differences. *n*_*m*_ represents the number of measurements in each split dataset.

| Dataset | Model | $n_m$ | Agreement Rate (Angular Diff.) | | AFD Diff. |
| --- | --- | --- | --- | --- | --- |
|  |  |  | Single Fiber | Two Fibers |  |
| dHCP | MSMT-CSD | 150 | 92.4 % (6°) | 44.2 % (16°) | 0.206 ± 0.08 |
| dHCP | SS3T-CSD | 54 | 62.7 % (6°) | 55.9 % (13°) | 0.066 ± 0.01 |
| BCP | MSMT-CSD | 75 | 86.0 % (6°) | 45.5 % (17°) | 0.117 ± 0.05 |

For AFD differences, SS3T-CSD achieves significantly lower error rates. For the different experiments, metrics related to voxels with 3-fibers are not reported because of their low consistency.

### 4.2 Intra-site experiments

#### 4.2.1 dHCP

We generally observe (Figure 3) similar performances of the different models for the agreement rate and the angular error when the number of input diffusion gradient directions is 15 or higher. For six directions, U-Net models outperform in both metrics (agreement rate and angular error) for both MSMT- and SS3T-CSD reference reconstructions serving as training GT. This can be explained by the large field of view (i.e., patches of 16^3^ of that method that compensate for the small number of directions. This trend is reversed for higher input directions where CTtrack and the MLP score are slightly higher, especially for SS3T-CSD.

Interestingly, and as can be depicted qualitatively in the left column of Figure 2 (shown for the case of U-Net), 2-fibers errors are significantly lower for SS3T-CSD than MSMT-CSD. This is consistent with the power of SS3T-CSD in depicting crossing fibers compared to MSMT-CSD in newborns (Dhollander et al., 2019). This can be observed in Figure 4, where crossing fibers between the cortico-spinal tract and the corpus callosum are clearly delineated for SS3T-CSD, where the gray matter component is not overestimated as in MSMT-CSD. For anatomical reference, we include the corresponding slice from the atlas by Pietsch et al. (2019).

**Figure 2.**
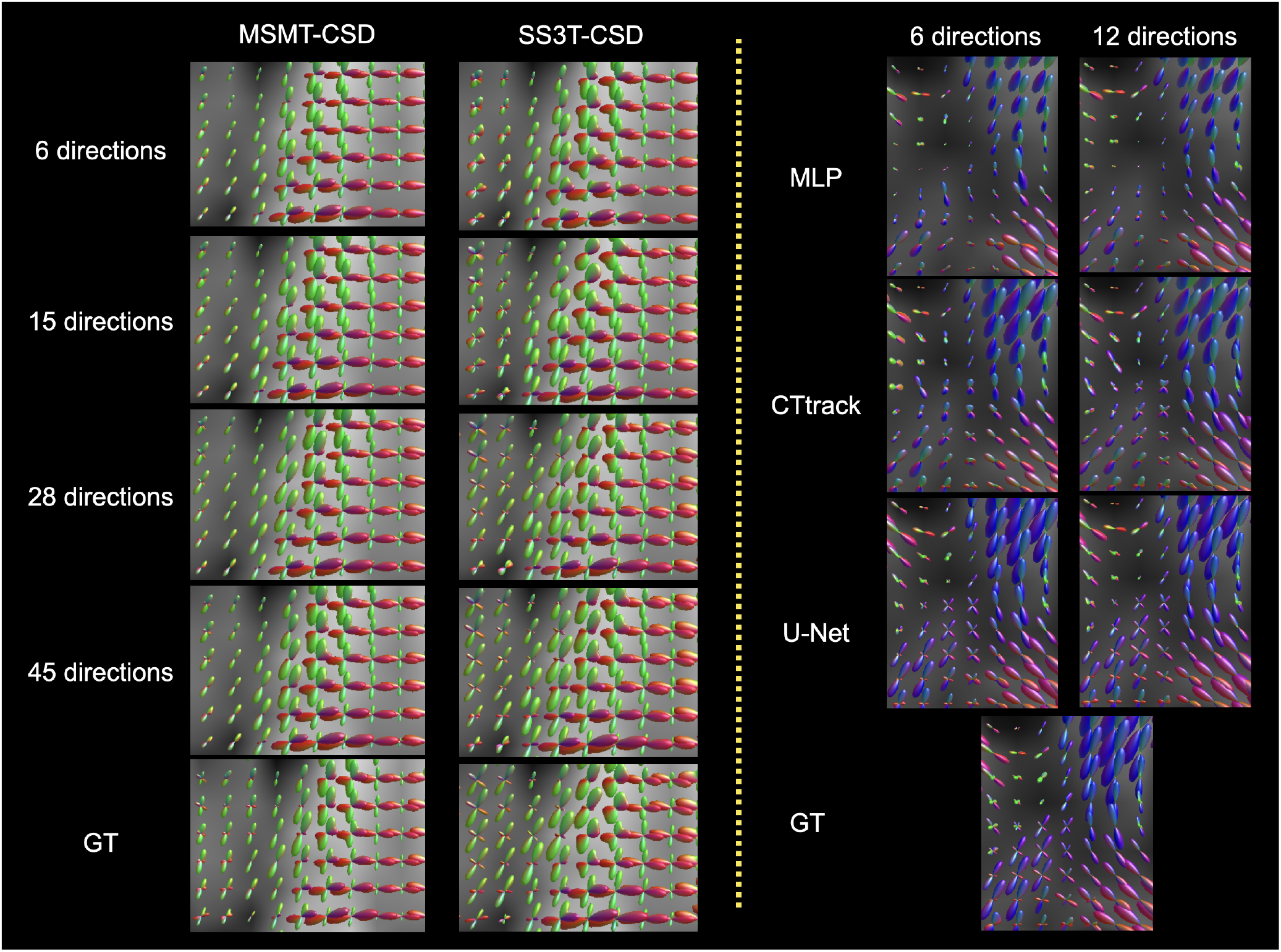
Qualitative examples of fODFs estimated for the dHCP dataset (left column) as predicted by U-Net; and for the BCP dataset (right column) as predicted by MLP, CTtrack, and U-Net for a different number of input measurements.

**Figure 3.**
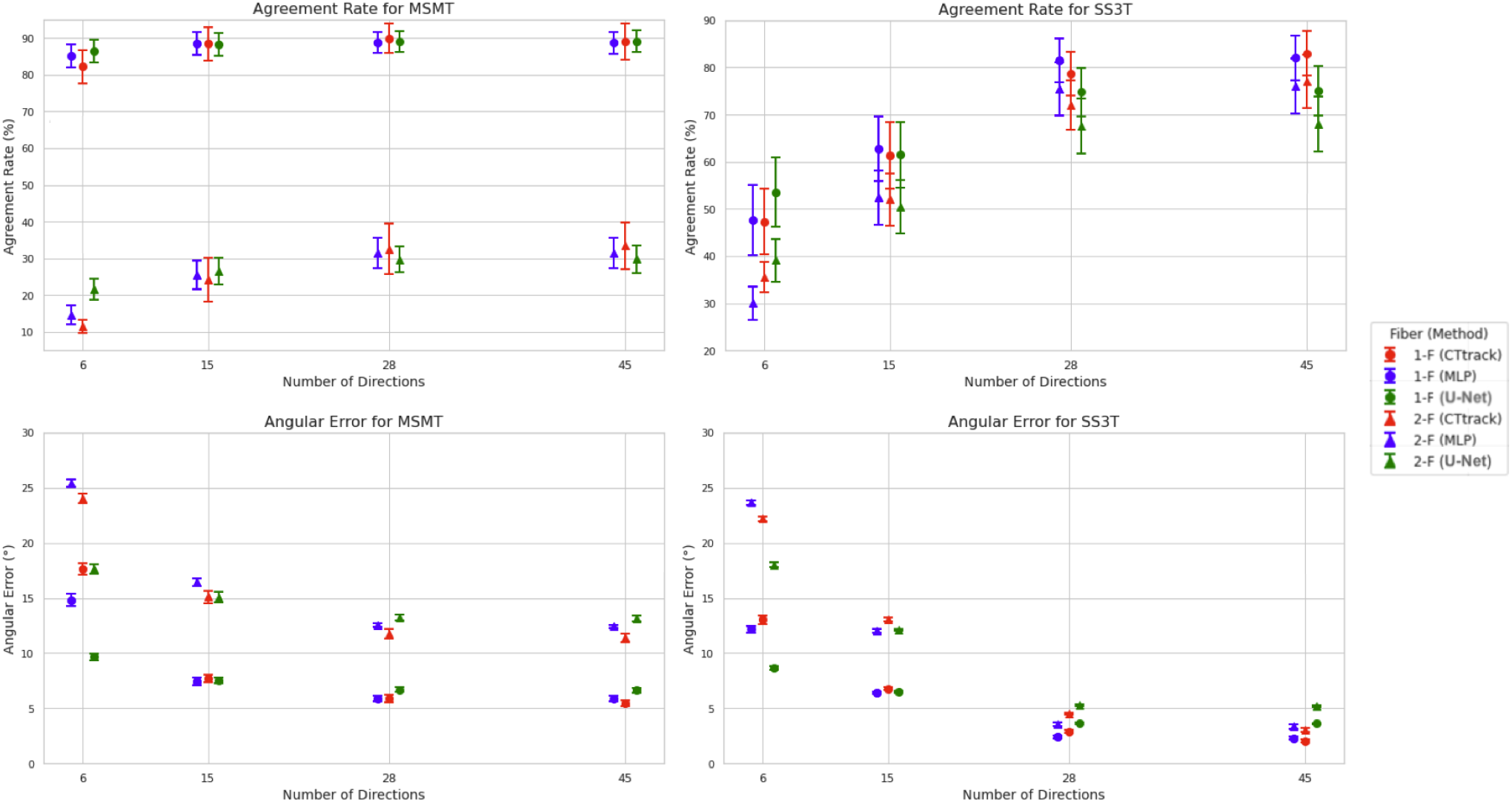
Model performances in terms of agreement rate (top row) and angular error (bottom raw) for both MSMT-CSD (left column) and SS3T-CSD (right column) trained and tested on the dHCP dataset. Metrics are reported for different numbers of input diffusion gradient directions and all methods (MLP, CTtrack, and U-Net).

**Figure 4.**
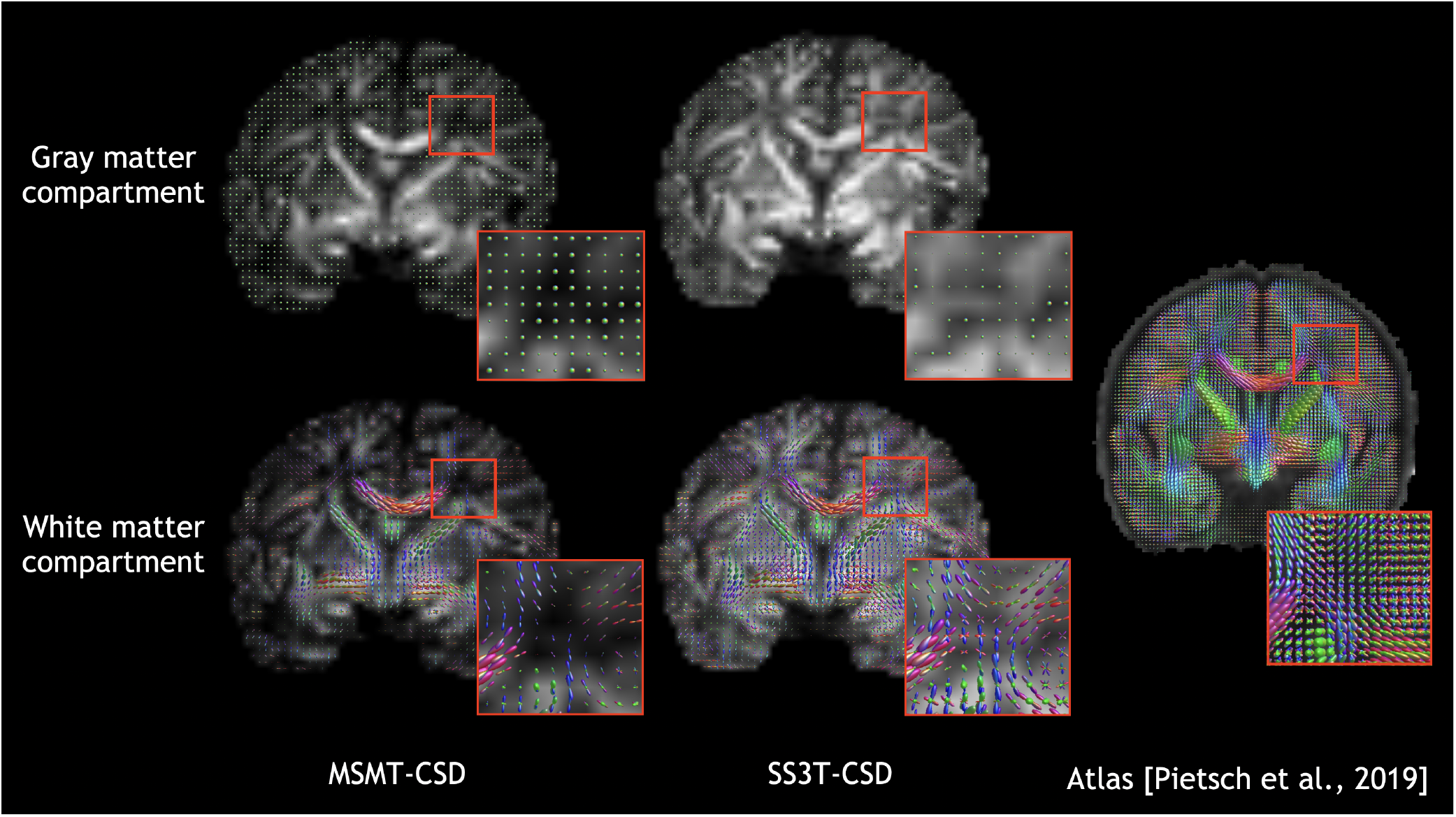
Qualitative comparison of coronal slices of the fODFs used as training GT for the learning-based models, reconstructed with MSMT-CSD and SS3T-CSD, respectively, from the dMRI scan of a subject at 40 weeks PMA. The visualization presents both gray and white matter compartments overlaid on fractional anisotropy (FA) maps derived from the dMRI data. The reconstructions were generated and visualized using MRtrix (Tournier et al., 2012) and its fork MRtrix3Tissue (https://3Tissue.github.io). For anatomical reference, the rightmost image shows the corresponding slice from the brain atlas published by Pietsch et al. (2019). Inset boxes highlight detailed regions of the reconstructions, demonstrating the fiber orientation patterns in both compartments.

We also observe a plateau in performance for all the methods from 28 to 45 directions. This plateau is close to being reached quickly with 15 directions for MSMT-CSD.

For the AFD error (Supplementary Figure 9, top row), we make similar observations as for AR and AE regarding the edge of U-Net compared to other models for six directions, especially for SS3T-CSD. However, this edge is kept for all input direction configurations.

#### 4.2.2 BCP

We observe (Figure 5) an overperformance of U-Net compared to the other methods that are more pronounced for 2-fibers, for both agreement rate in the number of peaks and their angular error, as also qualitatively shown in Figure 2 (right column). A similar observation can be made for the AFD error (Supplementary Figure 9, bottom row). Moreover, we do not notice any significant improvement, as in neonatal dHCP, by going with more directions (in this case, from 6 to 12). This can perhaps be explained by the consistency of the BCP data compared to dHCP, which has more variations in the growing anatomy due to their early age.

**Figure 5.**
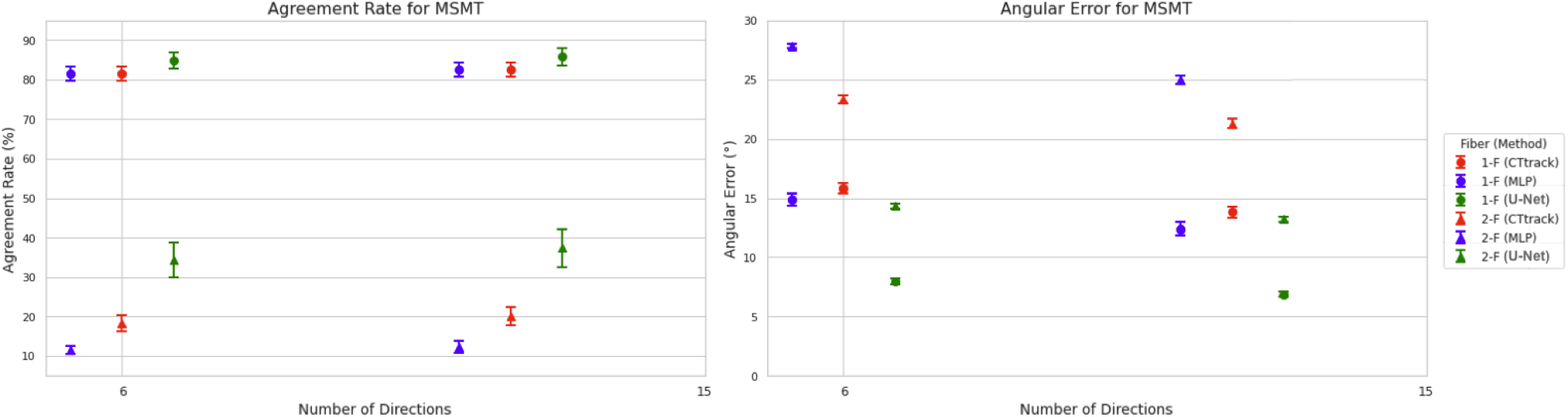
Model performances in terms of agreement rate (left column) and angular error (right column) the MSMT-CSD model trained and tested on the BCP dataset. Metrics are reported for different numbers of input directions (6 and 12) and all methods (MLP, CTtrack, and U-Net).

### 4.3 Age-related experiments in dHCP

From Figure 6a (AR) and Figure 6b (AE), we observe that when training in the late subset and tested in the early subset, domain shift is drastically reduced compared to the opposite. This can be observed especially for 2-fibers. Moreover, SS3T-CSD seems to be more robust in terms of the age domain shift compared to MSMT-CSD for both training on early or late subjects. Concerning the three models, they continue to score similarly across domain shifts, and U-Net continues to be the best model when training on 6 input directions and the opposite for higher directions. However, MLP is the only model that still improves slightly from 28 to 45 directions, especially in SS3T-CSD.

**Figure 6.**
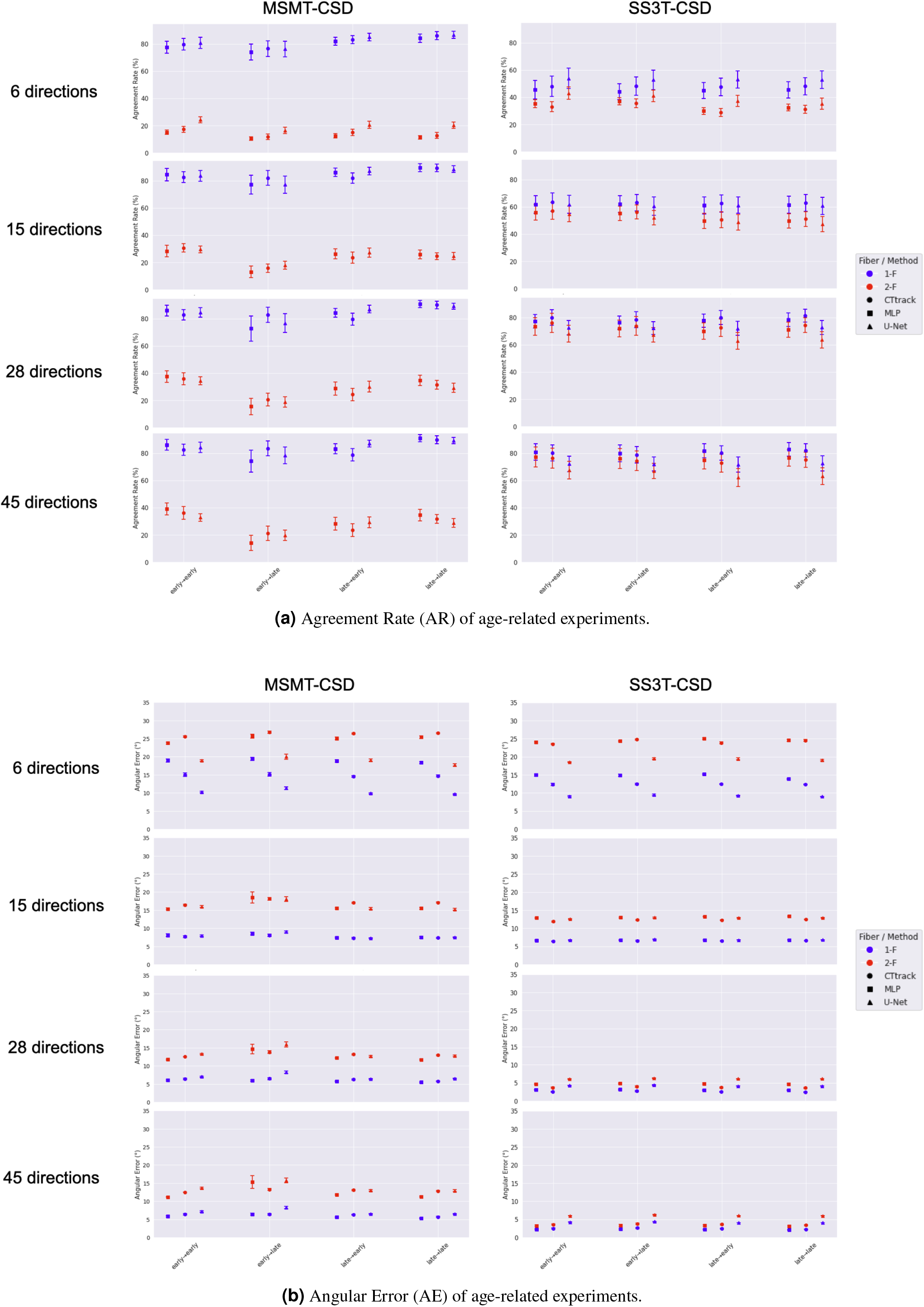
Age domain shift related experiments on the dHCP datasets of the different combinations of training and testing on early/late cohorts (as defined in Section 4.2). Results are reported for all three methods (MLP, CTtrack, and U-Net): (a) the agreement rate and (b) the angular error.

**Figure 7.**
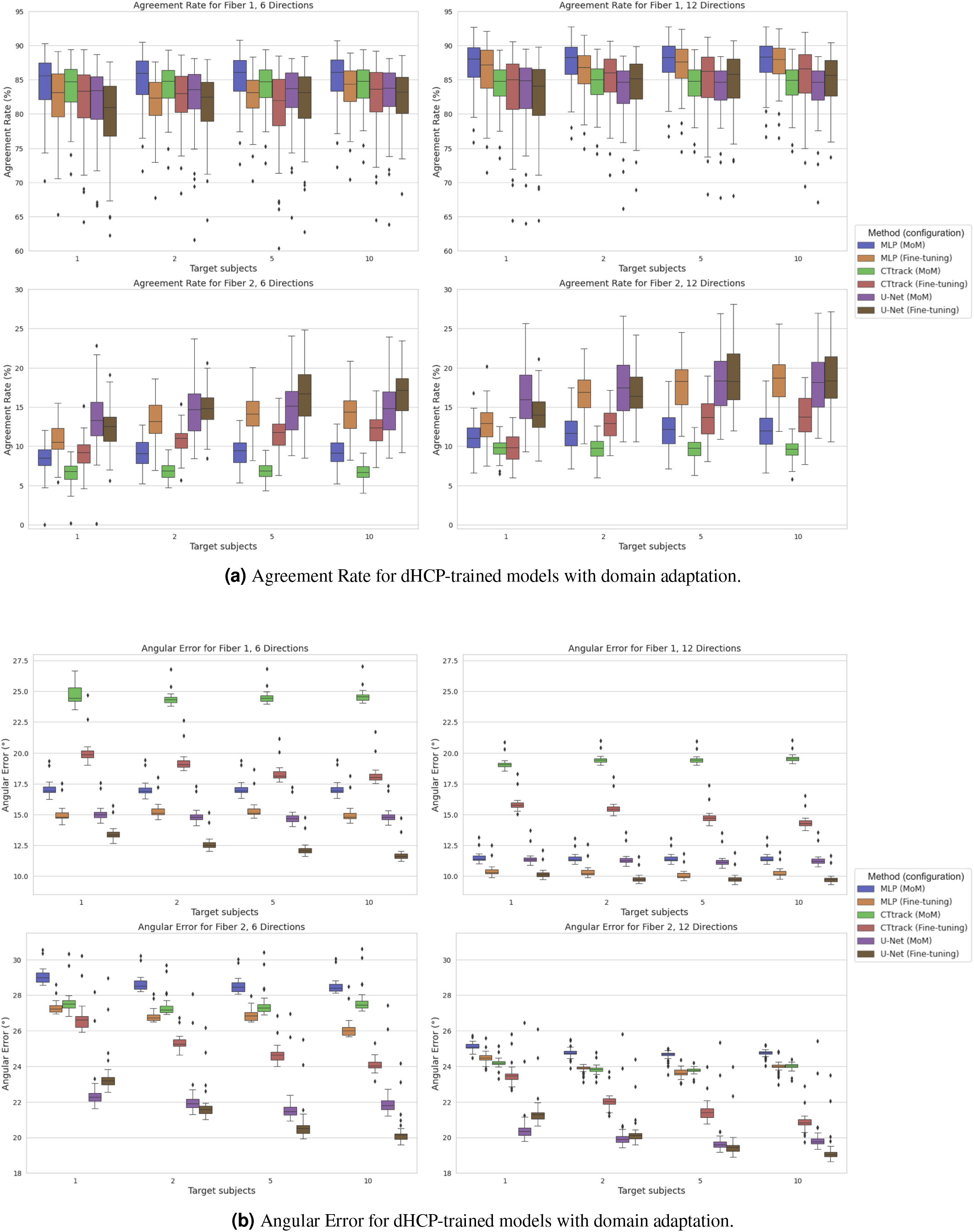
Performance metrics for dHCP-trained models with domain adaptation for target BCP testing, using a growing number of target domain subjects (1, 2, 5 and 10) for the different methods (MLP, CTtrack, and U-Net) using either the method of moments (MoM) or fine-tuning. Both (a) Agreement Rate (AR) and (b) Angular Error (AE) are reported for the case of 6 (left columns) and 12 (right columns) input directions.

**Figure 8.**
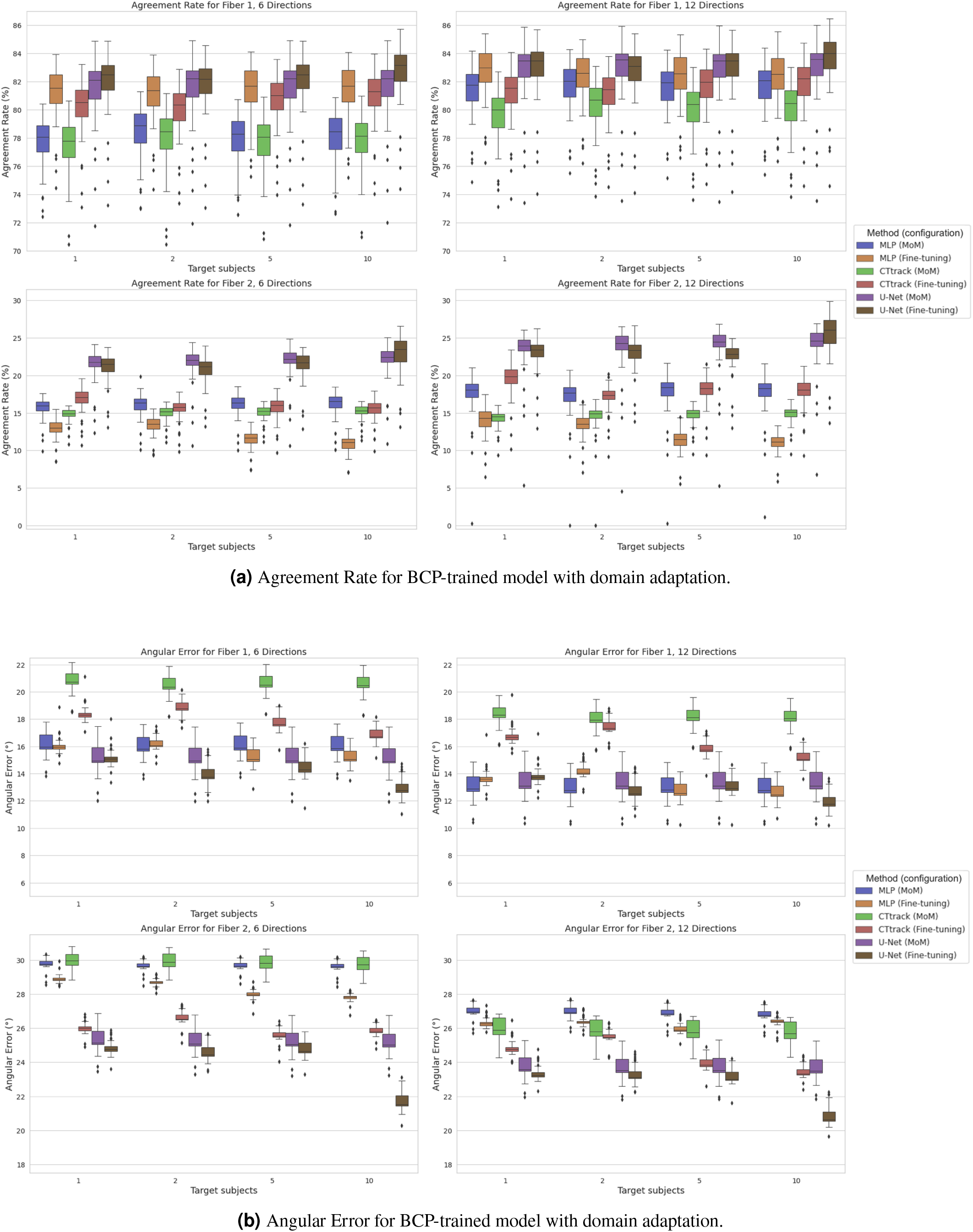
Performance metrics for BCP-trained models with domain adaptation for target dHCP testing, using a growing number of target domain subjects (1, 2, 5, and 10) for the different methods (MLP, CTtrack, and U-Net) using either the method of moments (MoM) or fine-tuning. Both (a) Agreement Rate (AR) and (b) Angular Error (AE) are reported for the case of 6 (left columns) and 12 (right columns) input directions.

For AFD (Supplementary Figure 10), U-Net still outperforms other methods in all configurations for six directions and in all MSMT-CSD cases when training on the late cohort and testing on the early one. The latter exhibits significantly higher errors compared to the other three age configurations. No significant difference, however, is observed across the other configurations, with domain shift being drastically reduced for SS3T-CSD as in the case of AR and AE.

### 4.4 Domain shift attenuation

#### 4.4.1 Trained on dHCP and tested on BCP

We generally observe that the single fiber configuration does not benefit as much as the 2-fibers configuration from an increased number of target subjects for domain shift attenuation for all models and both MoM and fine-tuning. Similarly, fine-tuning benefits more from the number of target subjects compared to MoM. Moreover, in the 2-fibers configuration, the more target subjects we add, the more fine-tuning overperforms MoM. For single fiber populations, MoM is slightly better or equal to fine-tuning in performance, especially for the angular error metric. Except single fiber configuration in agreement rate where MLP is overperforming other methods, we observe that U-Net is generally the best model in domain shift attenuation for both MoM and fine-tuning. When we go from 6 to 12 input diffusion gradient directions, MLP is the model that improves relatively more compared to other models.

For AFD error (Supplementary Figure 11-a), fine-tuning always surpasses MoM, with no significant improvement with the number of target subjects. This suggests that with a single subject, the distribution of the target AFD can be learned.

#### 4.4.2 Trained on BCP and tested on dHCP

We similarly observe that fine-tuning benefits more from increasing the number of target subjects compared to MoM for the angular error and the agreement rate. This can be particularly observed in AE. However, this increase is not very pronounced and only starts to show at 10 subjects. This can be particularly observed for the case of 2 fibers population. Fine-tuning generally outperforms the MoM, but not in all cases. For instance, for AR and 2-fiber populations, MLP is generally better with MoM compared to fine-tuning, and U-Net is better for the case of the number of target subjects if five or less. U-Net is generally the best model for addressing domain shifts, and it improves more with the number of target subjects and less with the number of gradient directions, the opposite of MLP and CTtrack.

For AFD error (Supplementary Figure 11-b), fine-tuning always surpasses MoM, with no significant improvement with the number of target subjects, up to 5 subjects as for AR and AE.

## 5 Discussion

In this study, we extensively investigated the performance and robustness of different deep-learning models on diffusion MRI-derived fODFs. We conducted intra-site experiments on two datasets, the developing Human Connectome Project (dHCP) and the Baby Connectome Project (BCP), and inter-site experiments to evaluate domain shift attenuation techniques. Specifically, we examined the effect of the number of input diffusion gradient directions, the influence of different ground truth configurations (MSMT- and SS3T-CSD), and age domain shift on model performance and employed two domain adaptation strategies: the Method of Moments (MoM) and fine-tuning.

Our intra-site experiments revealed that U-Net consistently outperformed the other models when fewer diffusion directions were used, particularly with the dHCP dataset and the SS3T-CSD ground truth. However, with an increased number of directions, MLP and CTtrack showed marginally but consistently better performance. Notably, SS3T-CSD exhibited lower angular errors for 2-fiber populations, highlighting its efficacy in capturing crossing fibers compared to MSMT-CSD in developing brains (Dhollander et al., 2019). This is also confirmed by the low consistency of the intra-ground-truth experiment, namely crossing fiber populations for the MSMT-CSD for which the agreement rate in the number of fibers is only 45%. The intra-site experiments also show a plateau in the performance observed across models from 28 to 45 directions, suggesting diminishing returns beyond a certain threshold of input directions and hence, a reduction in scanning time compared to non-deep-learning models. For SS3T-CSD, however, this performance is acceptable for both single and two-fibers, reaching around 75% in agreement rate and around 3% in the angular error.

In the BCP dataset, U-Net maintained its edge, particularly for 2-fiber populations, with less noticeable improvement when increasing from 6 to 12 directions. This could be attributed to the more consistent anatomical structures in older infants compared to the neonatal cohort of dHCP. Age-related experiments in dHCP demonstrated reduced domain shifts when training on later age groups, with SS3T-CSD proving more robust to age-related variability than MSMT-CSD.

For domain shift attenuation, fine-tuning generally surpassed MoM, particularly as the number of target subjects increased. U-Net emerged as the most reliable model for domain adaptation across different scenarios, with its performance benefiting more from an increase in target subjects than from additional gradient directions. For most of the configurations, fine-tuning with five target subjects yielded satisfactory results, as also demonstrated in other works such as semantic segmentation (Lhermitte et al., 2024; Zalevskyi et al., 2024). This, however, depends on the consistency and variability of the few target subjects. Future work can focus on optimizing the choice of these target subjects and how synthetic data can help compensate for real data.

The performance of DL-based fODF estimation models is bounded by the quality and consistency of the data, in particular crossing fibers voxels. Challenges faced by current diffusion MRI models, including MSMT-CSD and SS3T-CSD, in accurately estimating multiple fiber populations and low angular crossing fibers have been shown by Schilling et al. (2018), and also in our algorithm consistency analysis comparison experiment. In fact, the performance saturation at around 28 input directions is likely not a neural network limit but a model-data limit. The more diverse and consistent the data, the more crossing fibers, and crossing angles can be resolved, even for as few as six directions, as shown in our experiments, bypassing theoretical limits.

It is important to note that obtaining absolute *ground truth* fiber orientations remains an open challenge in the field, as it would require extensive histological validation, which is rarely available, especially in developing brains. Classical reconstruction methods (Dhollander and Connelly, 2016; Jeurissen et al., 2014a; Tournier et al., 2004) serve as practical reference standards, despite being approximations of the underlying anatomy. These methods, based on physical models and mathematical constraints, provide reasonable estimates that have been validated through various indirect means, including anatomical studies and phantom experiments. Future work can also attempt to merge ground-truth models in a way that leverages each of their advantages.

Domain adaptation is a promising strategy to foster model generalization in medical imaging, and many methods have been proposed recently (Guan and Liu, 2021) and increase the likelihood of potential deployment in clinical settings. Some of these challenges in fODF estimation, linked to the spatial resolution, for example, have been attempted to be addressed recently by implicit neural representations (Dwedari et al., 2024), and others related to the broad heterogeneity of the input acquisition (gradient directions, b-values, scanner type, etc.) by Ewert et al. (2024) using an encoder-decoder framework to learn a latent representation of the signal and the b-vectors.

Overall, our study highlights the potential of deep learning for fODF estimation of developing brains while underscoring key challenges related to domain shifts. While domain adaptation techniques like MoM and fine-tuning offer promising solutions, further research is needed to refine these methods, particularly in selecting optimal target subjects and improving ground-truth models used to generate the GT fODFs. Ultimately, improving data consistency and model robustness will be crucial for translating these models into real-world clinical applications.

## 6 Acknowledgements

We gratefully acknowledge access to the facilities and expertise of the CIBM Center for Biomedical Imaging (Centre d’Imagerie BioMédicale), a Swiss research center of excellence founded and supported by Lausanne University Hospital (CHUV), University of Lausanne (UNIL), École Polytechnique Fédérale de Lausanne (EPFL), University of Geneva (UNIGE), Geneva University Hospitals (HUG), and the Leenaards and Jeantet Foundations.

This research was supported by grants from the Swiss National Science Foundation (grants 182602 and 215641); the U.S. National Institutes of Health, including awards from the National Institute of Neurological Disorders and Stroke (R01NS128281) and the Eunice Kennedy Shriver National Institute of Child Health and Human Development (R01HD110772); and the National Natural Science Foundation of China (grant 62472315). The views and opinions expressed in this work are solely those of the authors and do not necessarily reflect the official policy or position of the funding agencies.

The authors would like to thank Dr. Khoi Minh Huynh at the University of North Carolina at Chapel Hill for discussion and code sharing on the Method of Moments, and Hakim Ouaalam at Boston Children’s Hospital and Harvard Medical School for preprocessing BCP T2-weighted images. We also thank Dr. Erick J. Canales-Rodriguez at EPFL, CIBM, and CHUV, and Dr. Yasser Alemán-Gómez at CHUV for their support and discussion during the preparation phase of this work. Finally, we thank Anmin Liu at Tongji University for his logistical support.

This retrospective research study used open-source human subject data from the Developing Human Connectome Project and the Baby Connectome Project, respectively, where ethical approval was **not** required per the data licenses.

## 7 Supplementary Material

**Figure 9.**
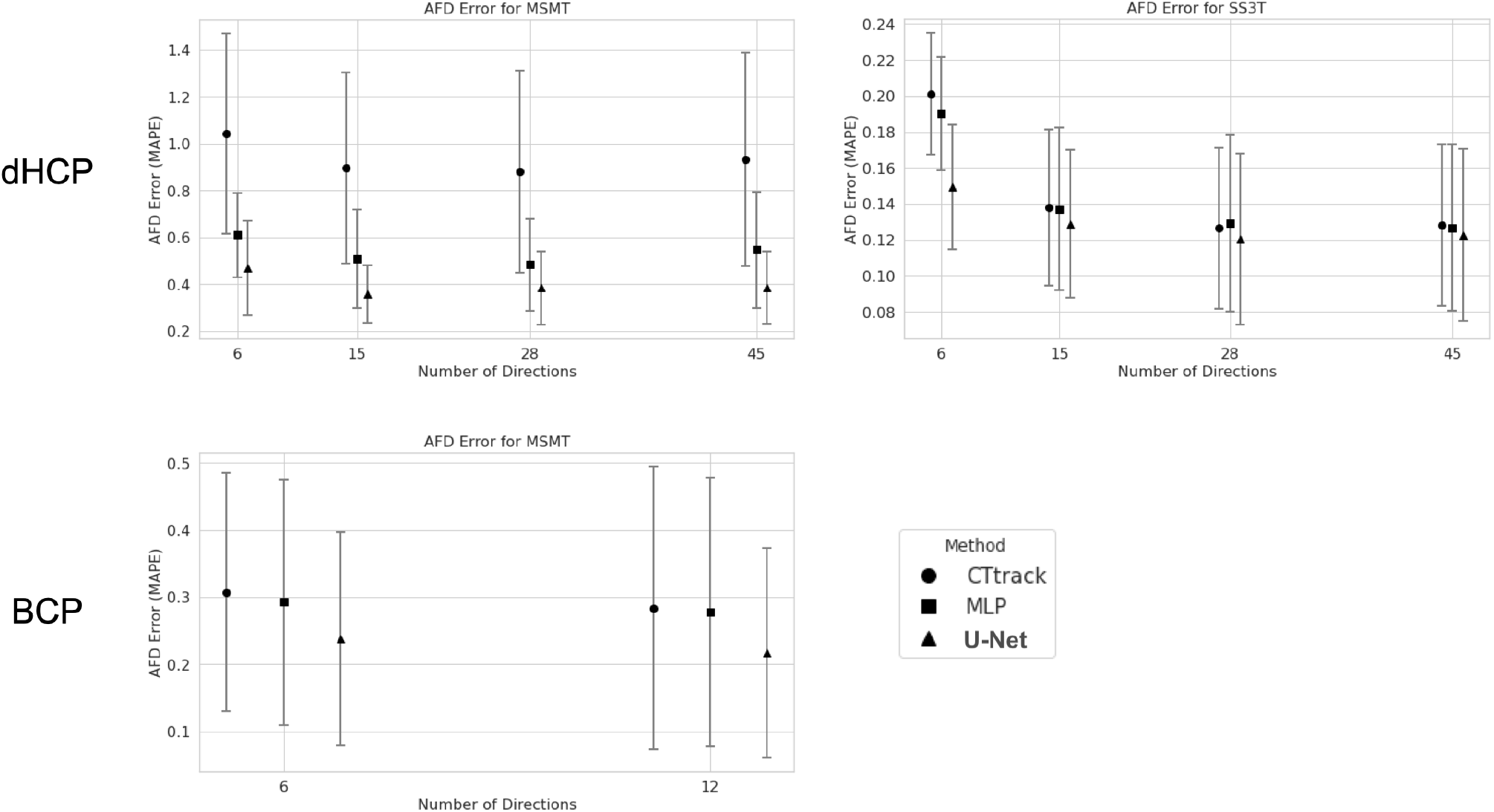
Model performances in terms of AFD error the MSMT-CSD model trained and tested on the dHCP (top row) and BCP (bottom row) datasets. Metrics are reported for different numbers of input directions and all three methods (MLP, CTtrack, and U-Net).

**Figure 10.**
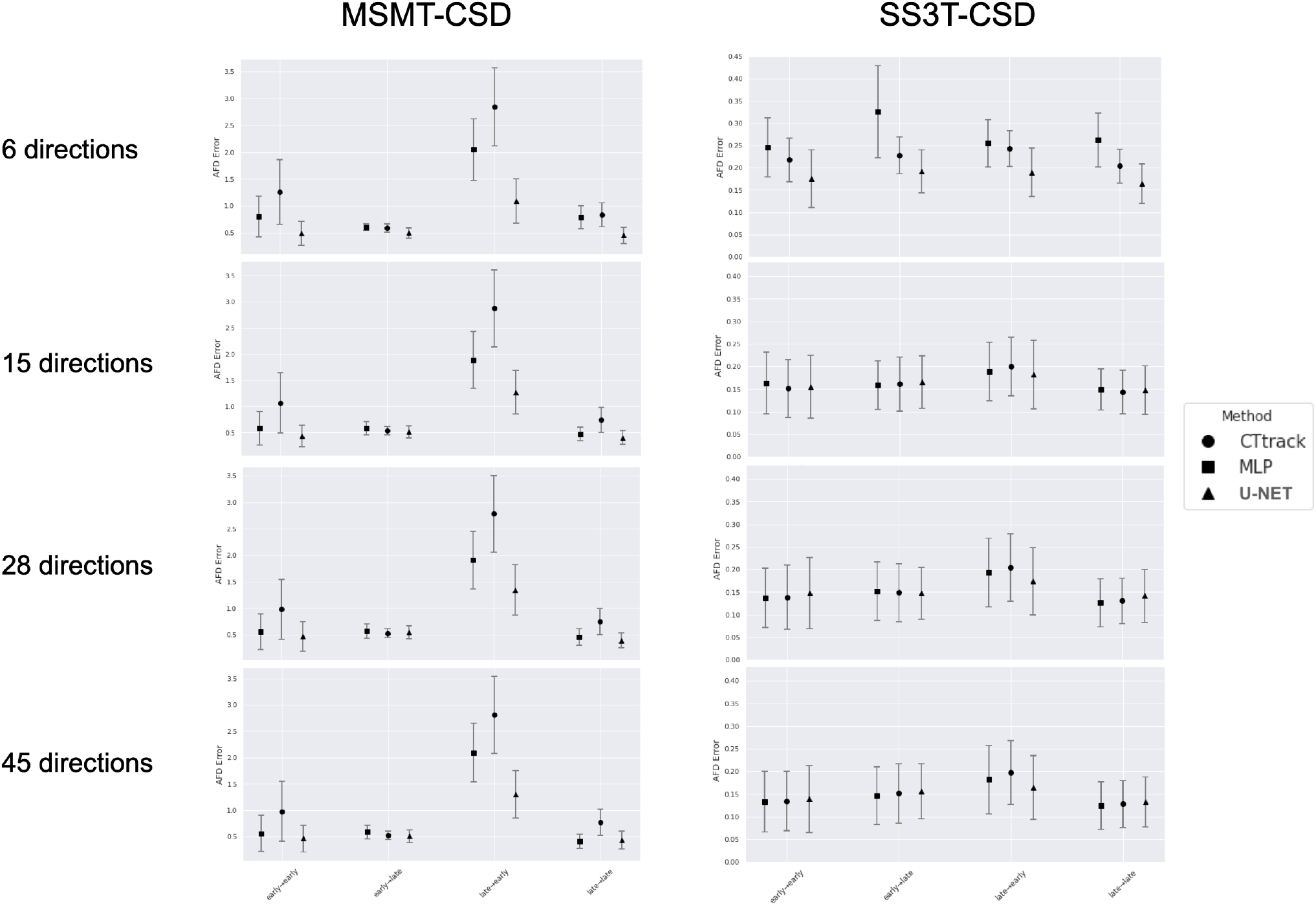
Age domain shift related experiments on the dHCP datasets of the different combinations of training and testing on early/late cohorts (as defined in Section 4.2). Results for the AFD error are reported for all three methods (MLP, CTtrack, and U-Net).

**Figure 11.**
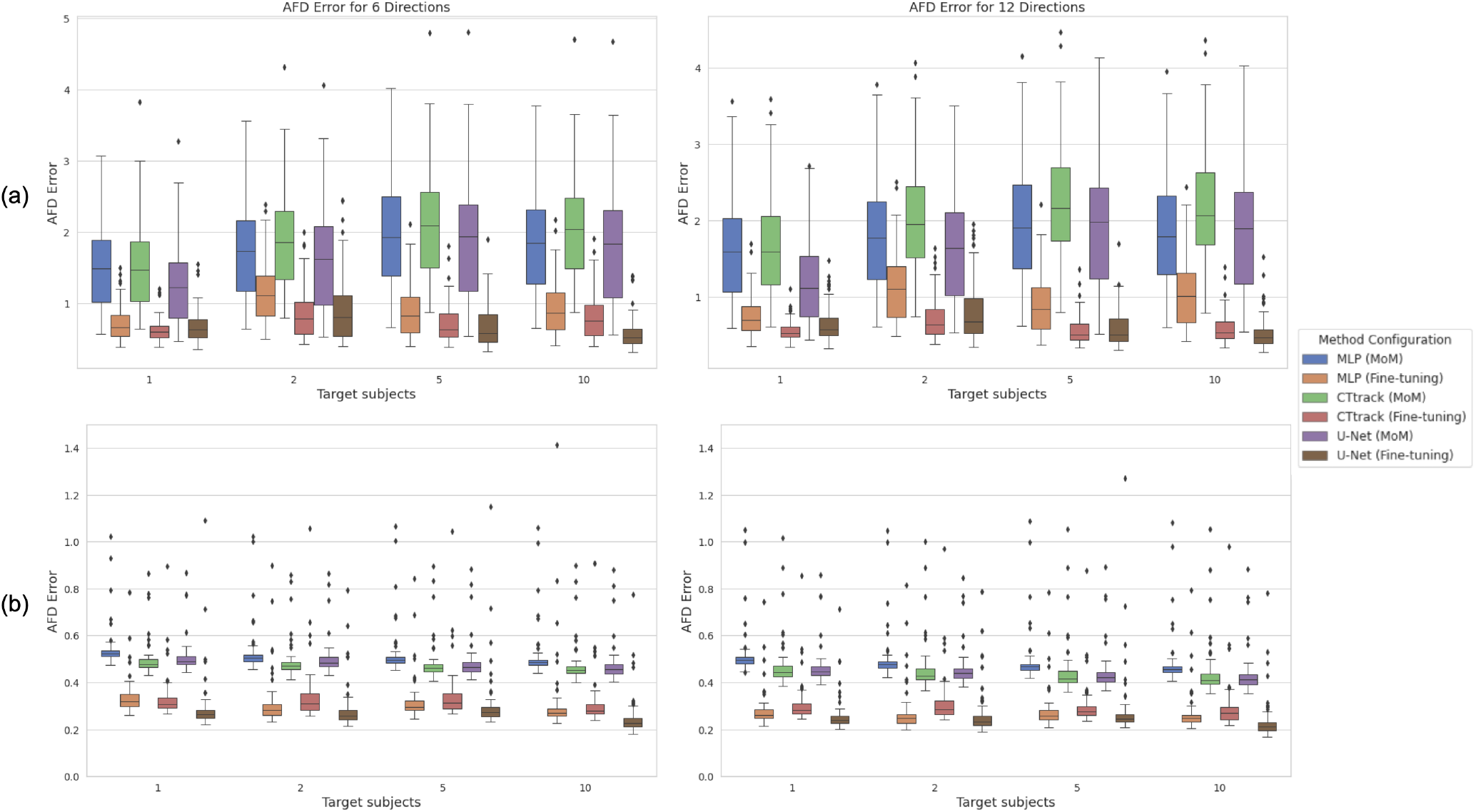
AFD error for (a) dHCP-trained models and (b) BCP-trained models with domain adaptation for target cross-testing, using a growing number of target domain subjects (1, 2, 5 and 10) for the different methods (MLP, CTtrack and U-Net) using either the method of moments (MoM) or fine-tuning, for the case of 6 (left columns) and 12 (right columns) input directions.

## Footnotes

1 https://www.developingconnectome.org/data-release/third-data-release/

2 https://www.humanconnectome.org/study/lifespan-baby-connectome-project/overview

